# Multiplex PCR method for MinION and Illumina sequencing of Zika and other virus genomes directly from clinical samples

**DOI:** 10.1101/098913

**Authors:** Josh Quick, Nathan D Grubaugh, Steven T Pullan, Ingra M Claro, Andrew D Smith, Karthik Gangavarapu, Glenn Oliveira, Refugio Robles-Sikisaka, Thomas F Rogers, Nathan A Beutler, Dennis R Burton, Lia Laura Lewis-Ximenez, Jaqueline Goes de Jesus, Marta Giovanetti, Sarah Hill, Allison Black, Trevor Bedford, Miles W Carroll, Marcio Nunes, Luiz Carlos Alcantara, Ester C Sabino, Sally A Baylis, Nuno Faria, Matthew Loose, Jared T Simpson, Oliver G Pybus, Kristian G Andersen, Nicholas J Loman

**Affiliations:** Institute of Microbiology and Infection, School of Biosciences, University of Birmingham, Birmingham, UK.; The Scripps Research Institute, La Jolla, CA, USA.; Public Health England, National infection Service, Porton Down, Salisbury, UK.; Department of Infectious Disease and Institute of Tropical Medicine, University of Saõ Paulo, Saõ Paulo, Brazil.; Scripps Translational Science Institute, La Jolla, CA, USA.; Massachusetts General Hospital, Boston, USA.; Instituto Oswaldo Cruz, Fundação Oswaldo Cruz, Rio de Janeiro; Fiocruz Bahia, Salvador, Brazil.; University of Rome, Tor Vergata.; Department of Zoology, University of Oxford, Oxford, UK.; Vaccine and Infectious Disease Division, Fred Hutchinson Cancer Research Center, Seattle, USA.; Department of Epidemiology, University of Washington, Seattle, USA.; University of Southampton, South General Hospital, Southampton, UK.; Instituto Evandro Chagas, Belem, Brazil.; Paul-Ehrlich-Institut, Langen, Germany.; University of Nottingham, Nottingham, UK.; OICR, Toronto, Canada.

**Author notes:** Correspondence should be addressed to N.J.L.

## Abstract

Genome sequencing has become a powerful tool for studying emerging infectious diseases; however, genome sequencing directly from clinical samples without isolation remains challenging for viruses such as Zika, where metagenomic sequencing methods may generate insufficient numbers of viral reads. Here we present a protocol for generating coding-sequence complete genomes comprising an online primer design tool, a novel multiplex PCR enrichment protocol, optimised library preparation methods for the portable MinION sequencer (Oxford Nanopore Technologies) and the Illumina range of instruments, and a bioinformatics pipeline for generating consensus sequences. The MinION protocol does not require an internet connection for analysis, making it suitable for field applications with limited connectivity. Our method relies on multiplex PCR for targeted enrichment of viral genomes from samples containing as few as 50 genome copies per reaction. Viral consensus sequences can be achieved starting with clinical samples in 1-2 days following a simple laboratory workflow. This method has been successfully used by several groups studying Zika virus evolution and is facilitating an understanding of the spread of the virus in the Americas.

## INTRODUCTION

Genome sequencing of viruses has been used to study the spread of disease in outbreaks ^1^. Real-time genomic surveillance is important in managing viral outbreaks, as it can provide insights into how viruses transmit, spread, and evolve ^1–4^. Such work depends on rapid sequencing of viral material directly from clinical samples. During the Ebola virus epidemic of 2013-2016, prospective viral genome sequencing was able to provide critical information on virus evolution and help inform epidemiological investigations ^3–6^. Sequencing directly from clinical samples is faster, less laborious, and more amenable to near-patient work than time consuming culture-based methods. Metagenomics, the process of sequencing the total nucleic acid content in a sample (typically cDNA or DNA), has been successfully applied to both virus discovery and diagnostics ^7–9^. Metagenomic approaches have seen rapid adoption over the past decade, fuelled by relentless improvements in the yield of high-throughput sequencing instruments ^5,10–12^. Whole-genome sequencing of Ebola virus directly from clinical samples without amplification was possible due to the extremely high virus copy numbers found in acute cases ^13,14^ However, direct metagenomic sequencing from clinical samples poses challenges with regards to sensitivity: genome coverage may be low or absent when attempting to sequence viruses that are present at low abundance in a sample with high levels of host nucleic acid background. During recent work on the Zika virus epidemic ^15^, we found that it was difficult to generate whole-genome sequences directly from clinical samples using metagenomic approaches (**Table 1**). These samples had Ct values between 33.9 and 35.9 (equivalent to 10-48 genome copies per microlitre). Before sequencing these samples were depleted of human rRNA and prepared for metagenomic sequencing on the Illumina MiSeq platform as previously described ^2,16^. In these cases, sequences from Zika virus comprised <0.01% of the dataset, resulting in incomplete coverage. Greater coverage and depth is critical for accurate genome reconstruction and subsequent phylogenetic inference. Additionally, there are significant sequencing and storage costs associated with generating large sequencing datasets, and these approaches currently do not lend themselves to lower-throughput portable sequencing devices such as the Oxford Nanopore MinION.

**Table 1.**
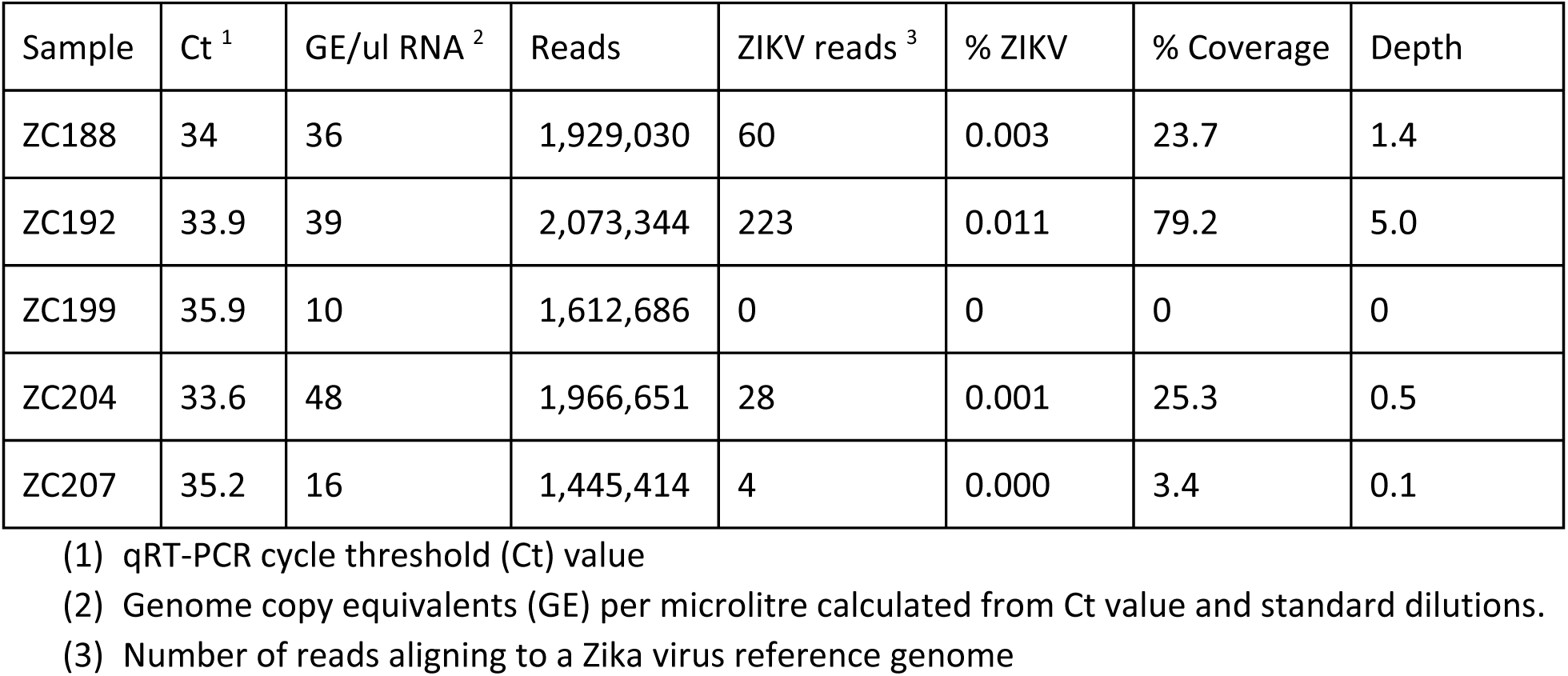
Results of metagenomic sequencing on five Zika-positive clinical samples collected from Columbia in January 2016

| Sample | Ct <sup>1</sup> | GE/ul RNA <sup>2</sup> | Reads | ZIKV reads <sup>3</sup> | % ZIKV | % Coverage | Depth |
| --- | --- | --- | --- | --- | --- | --- | --- |
| ZC188 | 34 | 36 | 1,929,030 | 60 | 0.003 | 23.7 | 1.4 |
| ZC192 | 33.9 | 39 | 2,073,344 | 223 | 0.011 | 79.2 | 5.0 |
| ZC199 | 35.9 | 10 | 1,612,686 | 0 | 0 | 0 | 0 |
| ZC204 | 33.6 | 48 | 1,966,651 | 28 | 0.001 | 25.3 | 0.5 |
| ZC207 | 35.2 | 16 | 1,445,414 | 4 | 0.000 | 3.4 | 0.1 |

(1) qRT-PCR cycle threshold (Ct) value
(2) Genome copy equivalents (GE) per microlitre calculated from Ct value and standard dilutions.
(3) Number of reads aligning to a Zika virus reference genome

Target enrichment is often required to generate complete viral genome coverage from clinical samples in an economic way. Enrichment can be achieved directly through i) isolation in culture, ii) oligonucleotide bait probes targeting the virus of interest, or iii) indirectly via host nucleic acid depletion. Amplification may also be required to generate sufficient material for sequencing (>5 ng for typical Illumina protocols, and 100-1000 ng for MinION). Polymerase chain reaction (PCR) can provide both target enrichment and amplification in a single step, and is relatively cheap, available, and fast compared to other methods. In order to generate coding-sequence complete coverage, a tiling amplicon scheme is commonly employed ^17–19^. During our work with Ebola virus we were able to reliably amplify >95% of the genome by sequencing eleven long amplicons (1-2.5 kb in length) on the MinION ^5^. The likelihood of long fragments being present in the sample reduces with lower virus abundance. Therefore, we anticipated that for viruses like Zika that are present at low abundance in clinical samples, we would be more likely to amplify shorter fragments. As an extreme example, a recent approach termed ‘jackhammering’ was used to amplify degraded HIV-1 samples stored for over 40 years and used 200-300 nucleotide (nt) amplicons to help maximise sequence recovery ^20^.

Using shorter amplicons necessitates a larger number of products to generate a tiling path across a target genome. To do this in individual reactions would require a large number of manual pipetting steps and therefore the potential for mistakes and a heightened risk of cross-contamination, as well as a greater cost in time and consumables. To solve these problems, we designed a multiplex assay to carry out hundreds of reactions in individual tubes. Our resulting step-by-step protocol allows any researcher to successfully amplify and sequence low abundance viruses directly from clinical samples.

### Description of the protocol

We describe a fully integrated end-to-end protocol for rapid sequencing of viral genomes directly from clinical samples. The protocol proceeds in four steps; i) multiplex primer pool design, ii) multiplex PCR protocol, iii) sequencing on MinION or Illumina instruments and iv) bioinformatics analysis and QC (**Figure 1**).

**Figure 1.**
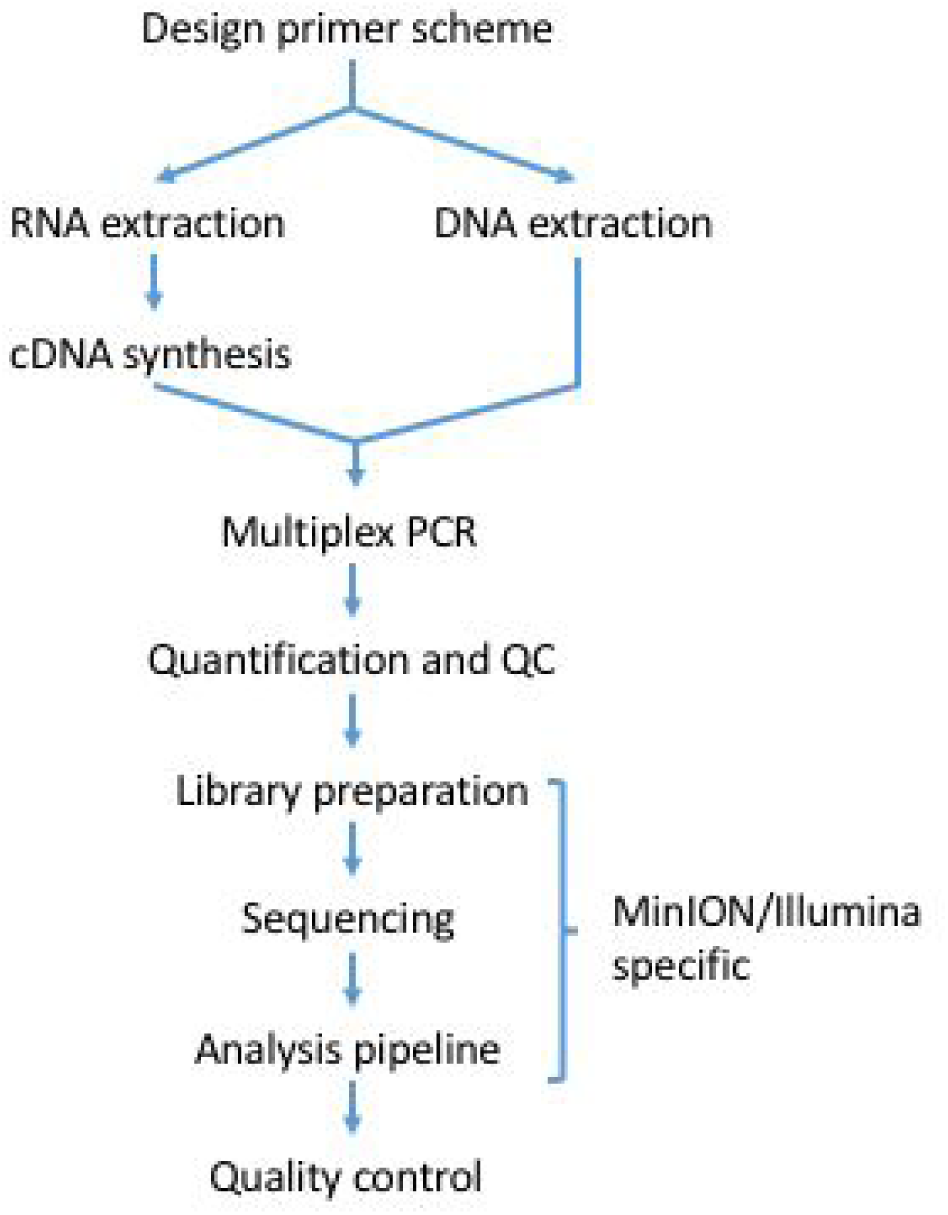
Workflow for tiling amplicon sequencing.

### Primer Design

We developed a web-based primer design tool entitled Primal Scheme (https://primal.zibraproject.org). which provides a complete pipeline for the development of efficient multiplex primer schemes. Each scheme is a set of oligonucleotide primer pairs that generate overlapping products. Together, the amplicons generated by the pairs span the target genome or region of interest (**Fig. 2**).

**Figure 2.**
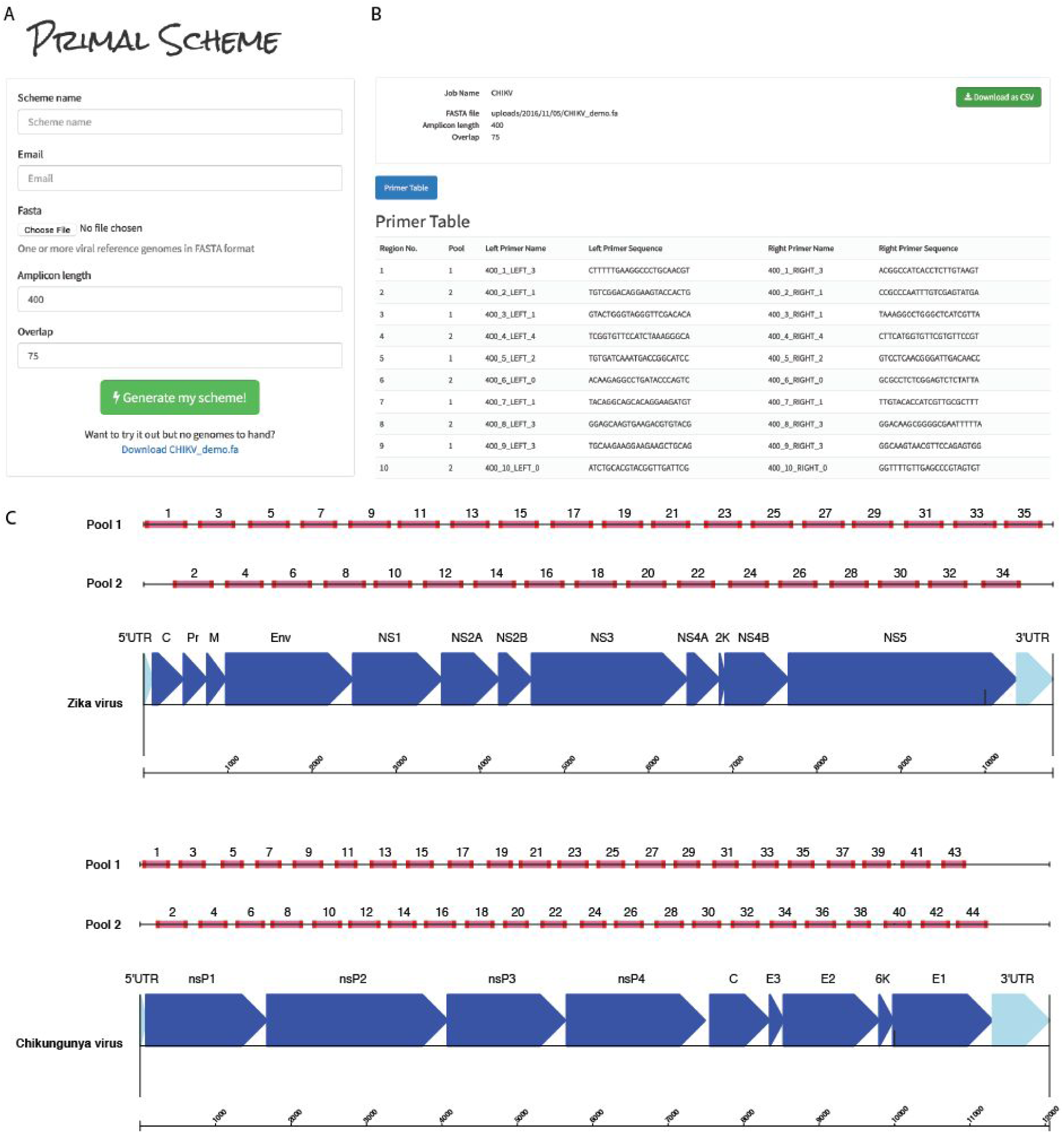
Overview of multiplex primer design using primal online primer design tool. (**a**) Submission box for online primer design tool. (**b**) Primer table of results. (**c**) Schematic showing expected amplicon products for each pool in genomic context for the ZikaAsian and ChikAsianECSA schemes.

As input, Primal Scheme requires a FASTA file containing one or more reference genomes. The user specifies a desired PCR amplicon length (default 400 nt, suggested values between 200 and 2000 nt), and the desired length of overlap between neighbouring amplicons (default 75 nt). Using a shorter amplicon length may be useful for samples where longer products fail to amplify (e.g. when the virus nucleic acid is highly degraded).

The Primal Scheme software performs the following steps:

- *Generate candidate primers:* The first sequence listed in the FASTA file should represent the most representative genome, with further sequences spanning the expected interhost diversity. Primal Scheme uses the Primer3 software to generate candidate primers ^21^. It chooses primers based on highly accurate thermodynamic modeling to take into account length, annealing temperature, %GC, 3’ stability, estimated secondary structure and likelihood of primer-dimer formation maximising the chance of a successful PCR reaction. Primers are designed with a high annealing temperature within a narrow range (65-68°C) which allows PCR to performed as a 2-step protocol (95°C denaturation, 65°C combined annealing and extension) for highly specific amplification from clinical samples without the need for nested primers.
- *Test candidate primers:* Subsequent reference genomes in the file are used to help choose primer pairs that maximise the likelihood of successful amplification for known virus diversity. A pairwise local alignment score between each candidate primer and reference is calculated to ensure the most ‘universal’ candidate primers are picked for the scheme. Mismatches at the 3’ end are severely penalised due as they have a disproportionate effect on the likelihood of successful extension^22,23^. The pairwise distances are summed, and the minimum scoring primer pairs are selected.
- *Output primer pairs:* Output files include a table of primer sequences to be ordered and BED file of primer locations used subsequently in the data analysis.

### Multiplex PCR Protocol

Next, we developed a multiplex PCR protocol employing novel reaction conditions, which allows amplification of products covering the whole genome in two reactions (**Fig. 3**). In comparison to single-plex methods, this dramatically reduces the cost of reagents and minimises potential sources of laboratory error. We assign alternate target genome regions to one of two primer pools, so that neighbouring amplicons do not overlap within the same pool (which would result in a short overlap product being generated preferentially). By screening reaction conditions based on concentration of cleaned-up PCR products, we determined that lower primer concentrations and longer annealing/extension time were optimal. Given the low-cost of the assay this step could also be performed alongside standard diagnostic qRT-PCR as a quality control measure to help reveal potential false positives ^24^.

**Figure 3.**
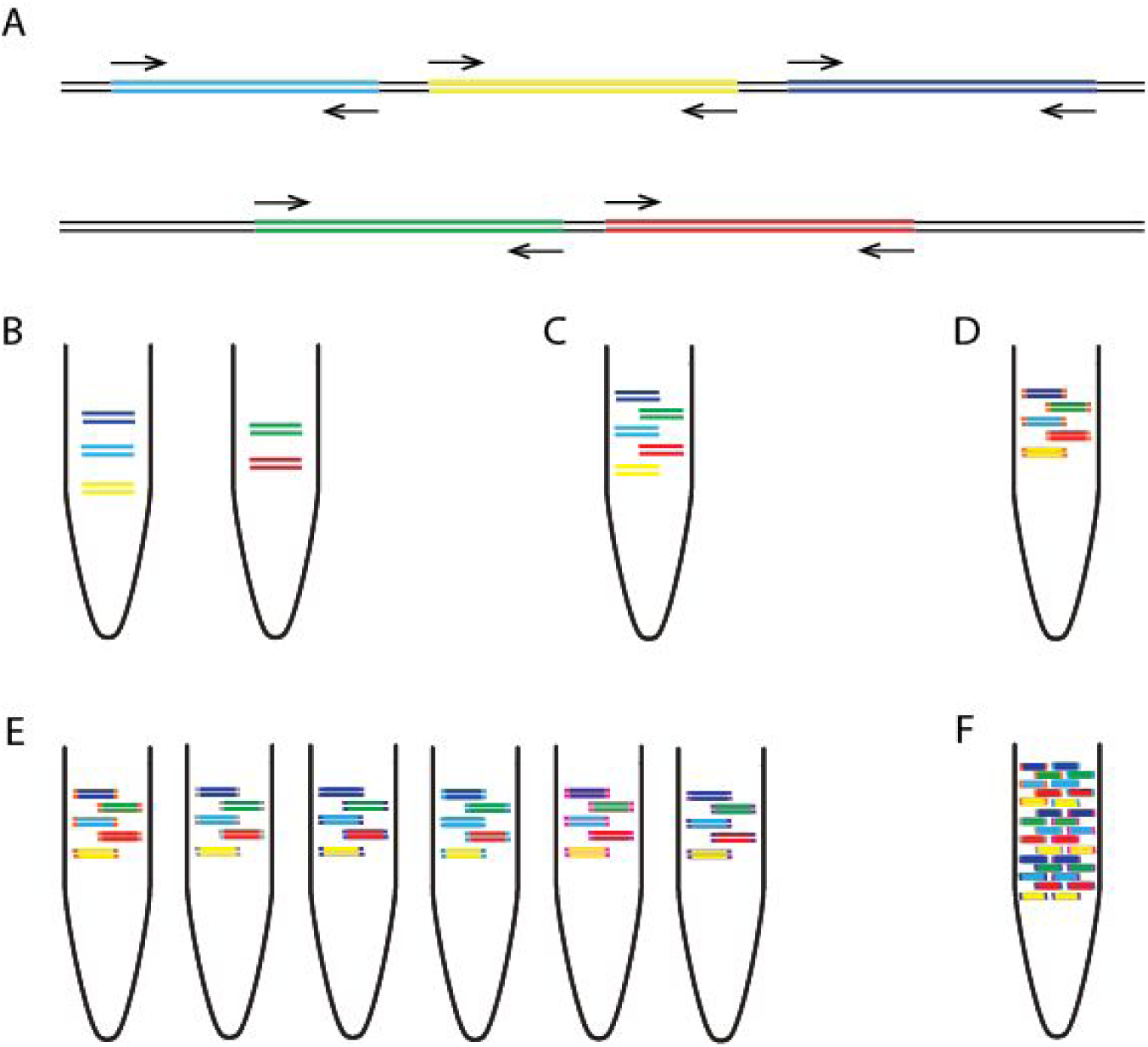
Overview of multiplex tiling PCR and pooling. (**a**) Schematic showing how the primers in pool ‘1’ and ‘2’ overlap between but not within reactions. (**b**) Amplicons generated by pool 1 and pool 2 primers from one sample are (**c**) pooled together, and (**d**) ligated with the same barcode during library preparation. (**e**) Samples are given unique barcodes so that (**f**) amplicons from many samples can be combined before platform specific sequencing.

### Sequencing protocol optimisations

Optimised library preparation methods for both the MinION and Illumina MiSeq platforms are provided, and should be readily adaptable to other sequencing platforms if required. The MinION system is preferred when portability and ease of setup in harsh environments is important ^5^. The Illumina platform is more suited to sequencing very large numbers of samples, due to greater sequence yields and the ability to barcode and accurately demultiplex hundreds of samples. Both platforms employ ligation-based methods to add the required sequencing adapters and barcodes. The molarity of adapters and cleanup conditions have been optimised for short amplicons. For the MinION we utilised the native barcoding kit (Oxford Nanopore Technologies, Oxford, UK) to allow up to 12 samples to be sequenced per flowcell. For the work presented here we used R9 or R9.4 flowcells (FLO-MIN105/FLO-MIN106) and the 2D library preparation kit (SQK-NSK007/SQK-LSK208). For the MiSeq platform the Agilent SureSelect^xt2^ adaptors and Kapa Hyper library preparation kit was used and allowed up to 96 samples per MiSeq run. Other library prep kits (e.g. Illumina TruSeq) and dual-indexed adaptors could also be used on the MiSeq. Depending on the number of reads required, the number of samples multiplexed, and the performance of the flowcell, sequencing on the MinION can be run for anything from a few minutes up to 72 hours. Typically 2-4 hours of sequencing is sufficient for 12 samples. For the MiSeq we recommend using the 2 x 250 nucleotide (nt) read-length for 400 nt amplicons, which can be completed in 48 hours.

### Bioinformatics workflow

A bioinformatics pipeline consisting of primer-trimming, alignment, variant calling, and consensus generation is made available for both Oxford Nanopore and Illumina platforms, the former building upon tools previously developed for Ebola virus sequencing in Guinea. The MinION pipeline is freely-available with components developed under the permissive MIT open source license at https://github.com/zibraproiect/zika-pipeline. The pipeline runs under the Linux operating system and is available as a Docker image meaning that it can also be run on Mac and Windows operating systems. This MinION pipeline can process the data from raw reads to consensus sequences on the instrument laptop given the correct primer scheme as a BED file and sample sheet as required input files.

### MinION pipeline

FAST5 reads containing signal-level information may be basecalled online using Metrichor, or offline using Albacore or nanonet in 2D mode. Nanonet is an open-source recurrent neural network (RNN) basecaller developed by Oxford Nanopore Technologies (https://github.com/nanoporetech/nanonet). The resulting FASTA files are demultiplexed by a script demultiplex.py into separate FASTA files for each barcode, as specified in a config file. By default these are set to the barcodes NB01-12 from the native barcoding kit. Alternatively, the Metrichor online service may be used to basecall read files and demultiplex samples. Each file is then mapped to the reference genome using `bwa mem` using the `–x ont2d` flag and converted to BAM format using `samtools view`. Alignments are preprocessed using a script that performs primer trimming and coverage normalisation. Primer trimming is performed by reference to the expected coordinates of sequenced amplicons, and therefore requires no knowledge of the sequencing adaptor (**Fig. 3**). Signal-level events are aligned using `nanopolish eventalign` and variants are called using `nanopolish variants`. Low quality or low coverage variants are filtered out and consensus sequences are generated using a script `margin_cons.py.` Variant calls and frequencies can be visualised using `vcfextract.py` and `pdf_tree.py.`

### Illumina Pipeline

First, we use Trimmomatic ^25^ to remove primer sequences (first 22 nt from the 5’ end of the reads) and bases at both ends with Phred quality score <20. Reads are aligned to the genome of a Zika virus isolate from the Dominican Republic, 2016 (GenBank: KU853012) using Novoalign v3.04.04 (http://www.novocraft.com/support/download/). SAMtools is used to sort the aligned BAM files and to generate alignment statistics ^26^. The code and reference indexes for the pipeline can be found at https://github.com/andersen-lab/zika-pipeline. Snakemake is used as the workflow management system ^27^.

### Alignment-based consensus generation

We have used an alignment based consensus approach to generate genomes as opposed to *de novo* assembly. Although *de novo* assembly could in theory be used with this protocol, the use of a tiling amplicon scheme already assumes the viral genome is present in a particular fixed order. This assumption may be violated in the presence of large-scale recombination. Some *de novo* assemblers, such as SPAdes, employ a frequency-based error correction preprocessing stage, and this may result in primer sequences being artificially introduced into the reference if primer sequences are not removed in advance ^28^. Importantly, when we compared alignment to *de novo*-based analysis methods of our generated Zika virus genomes, we found that we always obtained the same consensus sequences.

### Limitations of tiling amplicon sequencing

Our method is not suitable for the discovery of new viruses or sequencing highly diverse or recombinant viruses as primer schemes are virus-specific. Amplicon sequencing is prone to coverage dropouts that may result in incomplete genome coverage, especially at lower abundances, and the loss of both the 5’ and 3’ regions that fall outside the outer primer binding positions. The majority of primers are expected to work even when pooled equimolar, meaning largely complete genomes can be recovered without optimisation. The chikungunya data shown in **Table 3** was generated without any optimisation. However to achieve coding-sequence complete genomes, problem primers may need to be replaced or concentration relative to other primers adjusted in an iterative manner. Full coverage should be achievable for the majority of samples, however coverage is expected to correlate with viral abundance (**Table 4**). Targeted methods are also highly sensitive to amplicon contamination from previous experiments. Extreme caution should be taken to keep pre-PCR areas, reagents, and equipment free of contaminating amplicons. We have not tested this scheme with viral genomes longer than 12 kb. It is plausible that as the number of primer pairs increases that competitive inhibition may decrease PCR efficiency.

## MATERIALS

### REAGENTS

#### Tiling amplicon generation

QIAamp Viral RNA Mini Kit (Qiagen, cat. no. 52906)

Random Hexamers (50 μM) (Thermo Fisher, cat. no. N8080127)

Protoscript II First Strand cDNA Synthesis Kit (NEB, cat. no. E6560)

Deoxynucleotide (dNTP) Solution Mix (NEB, cat. no. N0447)

Q5 Hot Start High-Fidelity DNA Polymerase (NEB, cat. no. M0493)

PCR primers are listed in **Supplementary Table 1 and Table 2** (Integrated DNA Technologies)

Agencourt AMPure XP (Beckman Coulter, cat. no. A63881)

Qubit dsDNA HS Assay Kit (Thermo Fisher, cat. no. Q32854)

HyPre Molecular Biology Grade Water (GE Life Sciences, cat. no. SH30538.01)

100% ethanol

#### MinION sequencing

SpotON Flow Cell Mk I (R9.0) (Oxford Nanopore Technologies, cat. no. FLO-MIN105)

Nanopore Sequencing Kit (R9) (Oxford Nanopore Technologies, cat. no. SQK-NSK007)

Native Barcoding Kit (Oxford Nanopore Technologies, cat. no. EXP-NBD002)

NEBNext Ultra II End-repair/dA-tailing Module (NEB, cat no. E7546)

NEB Blunt/TA Ligase Master Mix (NEB, cat no. M0367)

MyOne C1 Streptavidin beads (Thermo Fisher, cat. no. 65001)

#### MiSeq sequencing

KAPA Hyper Library Prep (Roche, cat. no. 07962363001)

SureSelect^xt2^ indexes, MSQ, 16 (Agilent, cat. no. G9622A)

MiSeq Reagent Kit v2 (500 cycle) (Illumina, cat. no MS-102-2003)

D1000 ScreenTape (Agilent, cat. no. 5067-5582)

D1000 Reagents (Agilent, cat. no. 5067-5583)

KAPA Library Quantification Kit for Illumina platforms (Roche, cat. no 07960140001)

#### EQUIPMENT

Filtered pipette tips

1.5 ml microcentrifuge tube (Eppendorf, cat. no. 0030 108.051)

0.2 ml strip tubes with attached caps (Thermo Fisher, cat. no AB2000)

UV spectrophotometer (Thermo Fisher NanoDrop 2000, cat. no. ND-2000)

96-well thermocycler (Applied Biosystems Veriti, cat. no. 4375786)

Benchtop microcentrifuge (Thermo Fisher mySPIN 6, cat. no. 75004061)

Benchtop heater/shaker (Eppendorf ThermoMixer C)

Magnetic rack (ThermoFisher DynaMag-2, cat. no. 12321D)

PCR cabinet or pre-PCR room

#### MinION sequencing

MinION (Oxford Nanopore Technologies cat. no. MinION Mk1B)

Laptop with solid state disk (SSD) drive

#### MiSeq sequencing

MiSeq (Illumina)

TapeStation 2200 (Agilent)

## PROCEDURE

### Design and ordering of primers TIMING 1 h

1. (Optional) Identify representative reference sequences (e.g. from public databases such as GenBank) and generate a primer scheme using Primal Scheme by visiting primal.zibraproject.org. Choose an amplicon length that is suitable for your sequencing platform, and likely viral copy number of your sample (e.g. 300-500 nt for Zika on Illumina and MinION).
2. Order the primers generated by the online tool or from the pre-designed schemes provided in the **Supplementary Table 1 and 2.** Order pre-diluted in TE to 100 μM to avoid manually resuspending large number of primers.

#### Extraction of nucleic acids and preparation of cDNA if required TIMING 1-2 h

3| Tiling amplification is a general technique which can be applied to DNA or cDNA generated from RNA by reverse transcription. Follow the appropriate instructions for your application.

A. **RNA extraction and preparation of cDNA for RNA viruses**
  i. Extract RNA from 200 μl of either serum, plasma or urine using the QIAamp Viral RNA Mini Kit according to the manufacturer’s instructions eluting in 50 μl EB buffer.
  ii. Measure the 260/230 ratio using a spectrophotometer. Pure RNA should have a 260/280 ratio of 2.0 and a 260/230 ratio of 2.0-2.2. **TROUBLESHOOTING**
  iii. Perform the following two steps in a hood or dedicated pre-PCR area. Wash all surfaces with 1% sodium hypochlorite solution and irradiate labware with UV light for at least 10 minutes. Sample negative controls for each primer pool should be included on each PCR run. **CRITICAL STEP**
  iv. Mix the following in a 0.2 ml tube. This denaturation step minimises secondary structure in the RNA prior to cDNA synthesis.

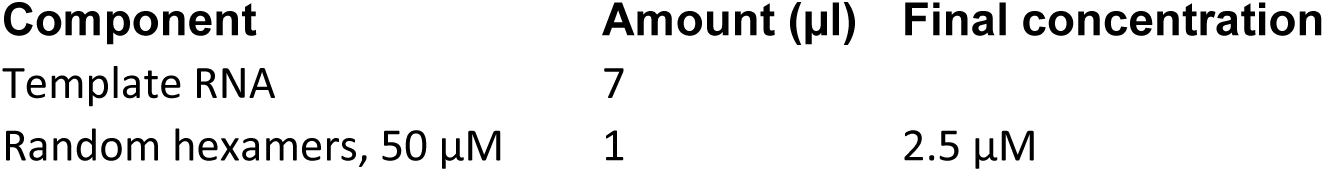

(v) Denature the template RNA by incubating on a heat block at 65 °C for 5 minutes before placing promptly on ice.
(vi) Complete the cDNA synthesis reaction by adding the following to the tube:

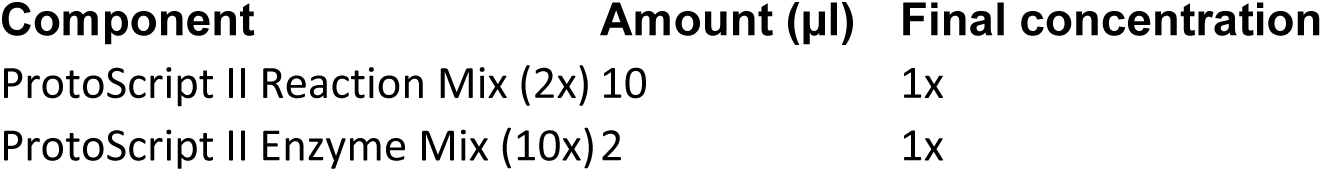

(vii) Perform the cDNA synthesis using the following conditions:

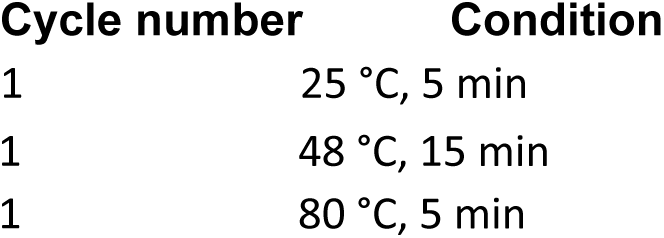

(B) **DNA extraction for DNA viruses**

i. Extract RNA from 200 μl of either serum, plasma or urine using the QIAamp MinElute Virus Spin Kit.
ii. Measure the 260/230 ratio using a spectrophotometer. Pure DNA should have a 260/280 ratio of 1.8 and a 260/230 ratio of 2.0-2.2. **TROUBLESHOOTING**

#### Preparing the primer pools TIMING 1 h

10 The first time the protocol is performed the primer pools must be prepared. Primers for alternate regions must be pooled so individual reactions overlap between pools but not within. 100 μM primers are combined into a stock primer pool that will be sufficient for amplifying many samples.
11 Label two 1.5 ml Eppendorf tubes using the scheme identifier and pool, either ‘1’ or ‘2’. Add an equal volume of each 100 μM primer stock such that both the forward and reverse primer for alternate regions are pooled together. For example. Pool ‘1’ would contain ZIKA_400_1_LEFT, ZIKA_400_1_RIGHT, ZIKA_400_3_LEFT, ZIKA_400_3_RIGHT, ZIKA_400_5_LEFT, ZIKA_400_5_RIGHT etc. for ZikaAsian scheme.

#### Performing multiplex tiling PCR TIMING 5 h

12 In the following step you will prepare the PCR reaction mixtures. A master mix of reagents is prepared for each pool as it reduces variability between reactions. The master mix should include some excess volume to allow for minor pipetting inaccuracy.
13 Prepare the following PCR reaction. The volumes given are for a 25 μl reactions but 50 μl can also be used. The volume of primers to use will depend on the number of primers in the pool as the final concentration should be 0.02 μM per primer. For example the ZikaAsian scheme from **Supplementary Table 1** has 36 primers in pool 1 so the volume to use would be 1.35 μl

| **Component** | **Amount (µl)** | **Final concentration** |
| --- | --- | --- |
| Q5 reaction buffer (5x) | 5 | 1x |
| dNTPs, 10 mM | 0.5 | 200 µM |
| Q5 DNA polymerase | 0.25 |  |
| PCR grade water | to 25 |  |
| Primer pool 1 or 2 (10 µM) | Varies | 0.015 µM per primer |
| cDNA or DNA | 2.5 |  |

14 On a thermocycler start the following program:

| **Cycle number** | **Denature** | **Anneal/extend** |
| --- | --- | --- |
| 1 | 98 °C, 30 s |  |
| 2-40 | 98 °C, 15 s | 65 °C, 5 m |

#### Cleanup and quantification of amplicons TIMING 1 h

15 Transfer contents of the tube to a 1.5 ml Eppendorf tube. Add the volume of SPRI beads given in the table below taking into account amplicon length. Complete clean-up according to the manufacturer’s instructions and elute in 30 μl nuclease-free water or EB.

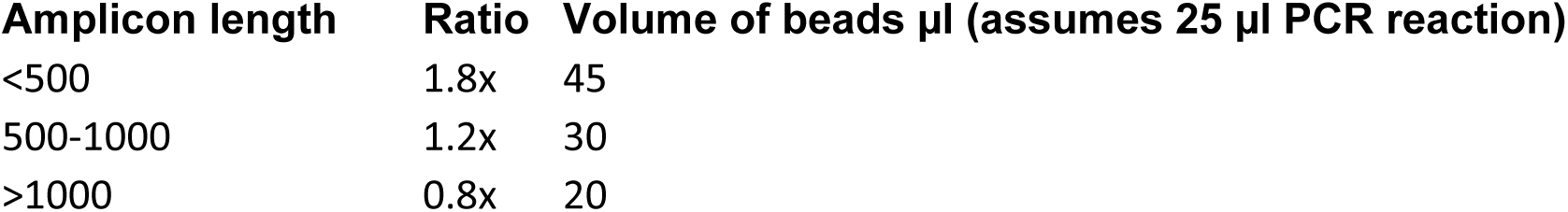

16 Quantify 1 μl of the cleaned product using the Qubit with the high-sensitivity assay as per the manufacturer’s instructions. You should expect concentrations in the range 5 - 50 ng/μl for each reaction from the Qubit quantification, except the PCR negative control which should be repeated if >1 ng/μl. **TROUBLESHOOTING**
17 (Optional) Mix 4 μl cleaned product and 1 μl 6x loading dye and run on a standard 1% agarose gel with a 1 Kb ladder. A specific band the correct size for your scheme should be observed on the gel. **TROUBLESHOOTING**

#### Library preparation and sequencing TIMING 24-28 h

18| Library preparation and sequencing are platform specific and and have been validated on the MinION from Oxford Nanopore Technologies and on the MiSeq from Illumina.

A. **Library preparation and sequencing using the MinION**
  i. *Number of samples per flowcell.* We recommend using two barcodes per sample (one barcode per pool per sample), which means that up to five samples and one negative control can be sequenced on each flowcell and allows each pool to be barcoded individually. This makes it easier to detect contamination that may be pool rather than sample specific. It also results in greater yield per sample, which improves genome coverage in samples with less even amplification.
  ii. *Normalisation.* To determine the correct quantity of amplicons to use per sample, divide 1500 ng by the number of non-negative control samples. For example, to sequence the recommended five samples for both primer pools (10 PCR reactions in total), 150 ng of amplicons from each PCR reaction would be used. PCR products should be kept separate at this stage, and the appropriate volume added to individual 1.5 ml Eppendorf tubes. The volume in each Eppendorf is then adjusted to 30 μl with nuclease-free water. **TROUBLESHOOTING**
  iii. *End-repair and dA-tailing.* Perform end-repair and cleanup according to the native barcoding protocol (Oxford Nanopore Technologies) eluting each in 15 μl nuclease-free water or EB.
  iv. *Barcode ligation.* In a 1.5 ml Eppendorf prepare the following ligation reactions, one reaction per amplicon pool.

| Component | Amount (µl) |
| --- | --- |
| dA-tailed amplicons | 15 |
| Native barcode NB01-NB12 | 5 |
| Blunt/TA Ligase Master Mix | 20 |

(v) Incubate at room temperature for 10 minutes followed by 65°C for 10 minutes to denature the ligase.
(vi) *Pool barcoded amplicons.* Combine all the barcode ligation reactions into a single 1.5 ml Eppendorf tube. Perform cleanup as before using an equal volume of SPRI beads and elute in 39 μl nuclease-free water. Since the pellet is large, you can speed up drying by briefly incubating the pellet at 50°C, checking intermittently to ensure the pellet does not overdry and crack.
(vii) *Sequencing adapter ligation.* In a 1.5 ml Eppendorf prepare the following ligation reaction. We have found using Blunt/TA Ligase instead of the NEBNext Quick Ligation Module as described in the native barcoding protocol improves the efficiency of this step.

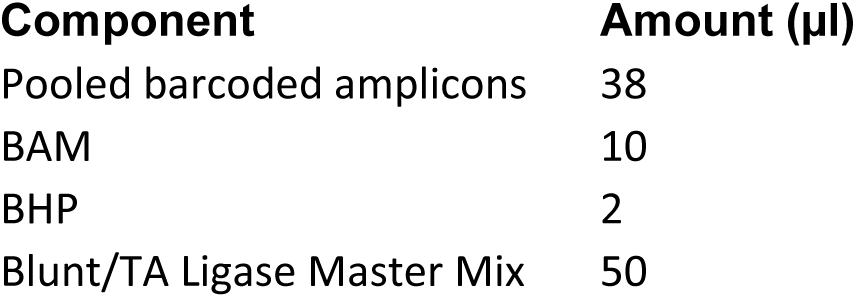

| Component | Amount (µl) |
| --- | --- |
| Pooled barcoded amplicons | 38 |
| BAM | 10 |
| BHP | 2 |
| Blunt/TA Ligase Master Mix | 50 |

(viii) *2D library preparation.* Complete library construction according to the native barcoding protocol. When purifying the adapted DNA with MyOne C1 beads, ensure that the pellet is fully resuspended after each wash, remove beads from the sides of the tube by scraping with a pipette tip if necessary. After the second wash, pipette off residual supernatant but do not leave the pellet dry. To prepare the library for loading, bring 100 ng of final library to a volume of 37.5 μl with nuclease-free water. This should be combined with 37.5 μl of RBF1. Excess library can be stored at 4°C for up to a week.
(ix) *Library loading.* Perform library loading and sequencing according to the manufacturer’s instructions. If the capillary action of the SpotON flowcell stops while loading library, restart the capillary action by loading another 200 μl of priming buffer through the inlet port then resume loading. **TROUBLESHOOTING**
(x) *Start sequencing run.* If you do not have an internet connection, ensure local basecalling is enabled by starting a script with a ‘_plus_Basecaller’ file name. Otherwise reads can be basecalled via Metrichor by starting the latest workflow for ‘2D Basecalling plus Barcoding’.

**(B) Library preparation, sequencing and analysis using the MiSeq.**

i. *Number of samples per flowcell.* We recommend using two barcodes per sample which means up to 47 samples plus a negative control can be sequenced on each run and allows each pool to be barcoded individually. This makes it easier to detect contamination that may be pool rather than sample specific. It also results in greater yield per sample which improves genome coverage in samples with more uneven amplification.
ii. *Normalisation.* Keeping pools in individual 1.5 ml Eppendorf tubes add 50 ng material and add nuclease-free water to adjust total volume to 50 μl.
iii. *End-repair and dA-tailing.* Perform end-repair and dA-tailing according to the Hyper prep kit protocol (KAPA).
iv. *Library preparation.* Complete library construction with the KAPA Hyper Prep Kit according to the manufacturer’s instructions substituting the KAPA Adapters for SureSelect^xt2^ Indexing Adaptors in the adapter ligation step. A 0.8X instead of 1X SPRI cleanup should be performed during the post-amplification cleanup to remove potential adaptor-dimers.
v. *Library quality control.* Measure the size distribution of the library using the TapeStation 2200 according the manufacturer’s instructions.
vi. *Library pooling.* Calculate the molarity of each library using the KAPA Library Quantification according to the manufacturer’s instructions and pool libraries in an equimolar fashion.
vii. *Library dilution and loading.* Prepare library for loading onto the MiSeq according to the manufacturer’s instructions.
viii. *Basecalling and demultiplexing.* Basecalling and demultiplexing will be performed automatically on the instrument if the sample sheet is provided when starting the run.

#### Data analysis TIMING 1-2 h

19| Download Docker application for Linux, Mac or Windows from (https://www.docker.com/products/overview). Run the installer to setup the docker tools on your machine. You should now be able to open a terminal window and run the command `docker--version` without getting an error. The source code of the pipeline is also available via The Zika analysis pipeline is available from https://github.com/zibraproject/zika-pipeline.

20| Download the Zika pipeline image from DockerHub by typing `docker pull zibra/zibra` into the terminal window.

21|The Zika pipeline is compatible with both MinION data and Illumina data yet there are some differences in the data handling required.

22| Start a Docker container with the following command. To test the container is working, run: `docker run -t -i zibra/zibra:latest`

Docker containers do not have access to the file system of the computer they run within by default. You will need to provide access to a local directory in order to see data files. This is achieved using the −v parameter. You may need to grant access to Docker to share the drive via the Shared Drives menu option under Settings. For example, on Windows, if you wished to provide access to the c:\data\reads directory to the Docker container, use the following `docker run -v c:/data/reads:/data -t -i zibra/zibra:latest`

Then, within the Docker container the /data directory will refer to `c:\data\reads` on the server.

23| Run the platform specific pipeline. The Zika pipeline is compatible with both MinION data and Illumina data yet there are some differences in the data handling required.

A. **Running analysis pipeline on MinION data.**
  i. Ensure the reads are basecalled using either Metrichor or an offline basecaller. Compatible basecallers including Albacore (available as installable packages for Linux, Windows and Mac through the MinION Community Portal) or and the freely-available and open-source nanonet (https://github.com/nanoporetech/nanonet) software. Nanonet is compatible with graphics processing unit (GPU) cards to increase speed.
  ii. Metrichor will perform demultiplexing if a barcoding workflow is selected. For other basecallers you must demultiplex reads manually using a script in the Docker image with the command: `demultiplex.sh <directory of FAST5 Files> <output directory>`
  iii. Run the Zika pipeline. The pipeline takes three required items `sample_id` -- the sample name (should not contain space characters) `directory` -- the directory containing the FAST5 files (e.g. output directory from previous step) `scheme` -- the name of the scheme directory, e.g. ZikaAsian `fast5_to_consensus.sh <sampleID> <directory> <scheme>` For example: `fast5_to_consensus.sh Zika1 /data/NB08/downloads/pass ZikaAsian`
  iv. Output files will be written to the current directory. The final consensus file will be named `<sampleID>.consensus.fasta`
B. **Running analysis pipeline on Illumina data.**
  i. Download and follow the instructions for the Illumina pipeline by reference to https://github.com/andersen-lab/zika-pipeline `illumina_pipeline.sh <sampleID> <fastq1> <fastq2> <scheme>`

#### Quality control TIMING 1 h

24 Check the coverage of the genomes by reference to the alignment file. Use an alignment viewer such as IGV^29^ or Tablet^30^ and load the `<sampleID>.primertrimmed.sorted.bam` file in conjunction with the reference sequence. Amplicons should be evenly spread throughout the genome. Deep piles of reads of single amplicons are potentially warning signs of contamination. The alignments should be compared with the negative control alignment to help indicate problematic samples or regions.
25 The Zika pipeline produces a variant frequency plot to allow you to rapidly determine the allele frequency of mutations in the sample (compared with the reference). The variant frequency plot is given the name `<sampleID>.variants.png` and is generated from the `<sampleID>.variants.tab` file that can be opened in spreadsheet applications or statistical software. The principle of the variant frequency plot is to identify mutations that occur at lower than expected allele frequencies and help decide whether they are a biological phenomenon (e.g. intra-host single nucleotide variants), potential signs of contamination or sequencing errors (for example in homopolymeric tracts in MinION data).

## TROUBLESHOOTING

Troubleshooting advice can be found in **Table 2.**

**Table 2.**
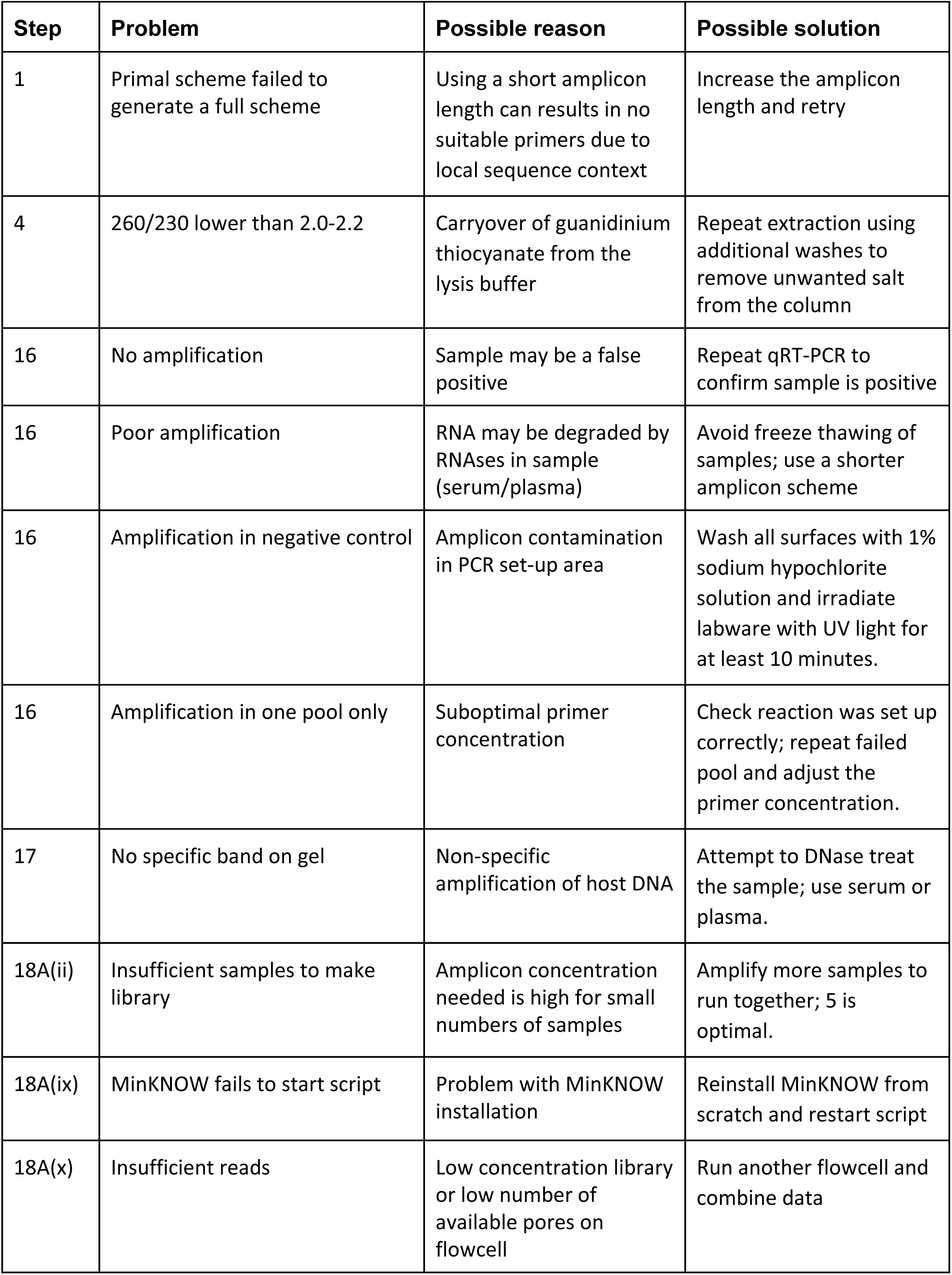
Troubleshooting table

| Step | Problem | Possible reason | Possible solution |
| --- | --- | --- | --- |
| 1 | Primal scheme failed to generate a full scheme | Using a short amplicon length can result in no suitable primers due to local sequence context | Increase the amplicon length and retry |
| 4 | 260/230 lower than 2.0-2.2 | Carryover of guanidinium thiocyanate from the lysis buffer | Repeat extraction using additional washes to remove unwanted salt from the column |
| 16 | No amplification | Sample may be a false positive | Repeat qRT-PCR to confirm sample is positive |
| 16 | Poor amplification | RNA may be degraded by RNases in sample (serum/plasma) | Avoid freeze thawing of samples; use a shorter amplicon scheme |
| 16 | Amplification in negative control | Amplicon contamination in PCR set-up area | Wash all surfaces with 1% sodium hypochlorite solution and irradiate labware with UV light for at least 10 minutes. |
| 16 | Amplification in one pool only | Suboptimal primer concentration | Check reaction was set up correctly; repeat failed pool and adjust the primer concentration. |
| 17 | No specific band on gel | Non-specific amplification of host DNA | Attempt to DNase treat the sample; use serum or plasma. |
| 18A(ii) | Insufficient samples to make library | Amplicon concentration needed is high for small numbers of samples | Amplify more samples to run together; 5 is optimal. |
| 18A(ix) | MinKNOW fails to start script | Problem with MinKNOW installation | Reinstall MinKNOW from scratch and restart script |
| 18A(x) | Insufficient reads | Low concentration library or low number of available pores on flowcell | Run another flowcell and combine data |

## TIMING

Steps 1-2, Design and ordering of primers: 1 h

Steps 3-4, RNA extraction and QC: 1 h

Steps 5-9, Preparation of cDNA: 1 h

Steps 10-11, Preparing the primer pools: 1 h

Steps 12-14, Performing multiplex tiling PCR: 5 h

Steps 15-17, Cleanup and quantification of amplicons: 1 h

Step 18, Library preparation and sequencing: 1-2 d

Steps 19-23, Data analysis: 1-2 h

Steps 24-25, Quality control of consensus sequences: 1 h

### ANTICIPATED RESULTS

This protocol should achieve near-complete genome coverage.

#### MinION Sequencing

As a demonstration of the ZikaAsian scheme on MinION we sequenced the World Health Organization Zika reference sample 11474/16^31^ (**Table 3**) and a chikungunya clinical sample from Brazil PEI-N11602. The Ct value for the Zika virus sample was between 18-20 depending on RNA extraction method used. The Ct value for the Chikungunya sample was determined by the RealStar^®^ Chikungunya RT-PCR Kit 1.0 from Altona Diagnostics (Hamburg, Germany). The Zika virus sample generated 97.7% coverage of the genome above 25x coverage. Coverage of the genome was reasonably even, with a drop-out in the middle of the genome (**Figure 4**). The WHO Control Reference MinION dataset is available from the CLIMB website (https://s3.climb.ac.uk/nanopore/Zika_Control_Material_R9.4_2D.tar).

**Table 3.** Results of MinION sequencing after barcode demultiplexing for an isolate of Zika and for a clinical sample of chikungunya virus.

| Sample | Scheme | Pore | Ct | # reads (2D pass) | Aligned reads | % aligned | % coverage | Depth |
| --- | --- | --- | --- | --- | --- | --- | --- | --- |
|  |  |  |  |  |  |  | (>25x) |  |
| Zika WHO Control Reference (11474/16) | ZikaAsian | R9.4 | 18-20 | 9,606 | 9,528 | 99.2 | 97.7 | 392 |
| Chikungunya PEI N11602 | ChikAsian nECSA | R9.4 | 20 | 89,365 | 88,527 | 99.1 | 88 | 2,779 |

**Figure 4.**
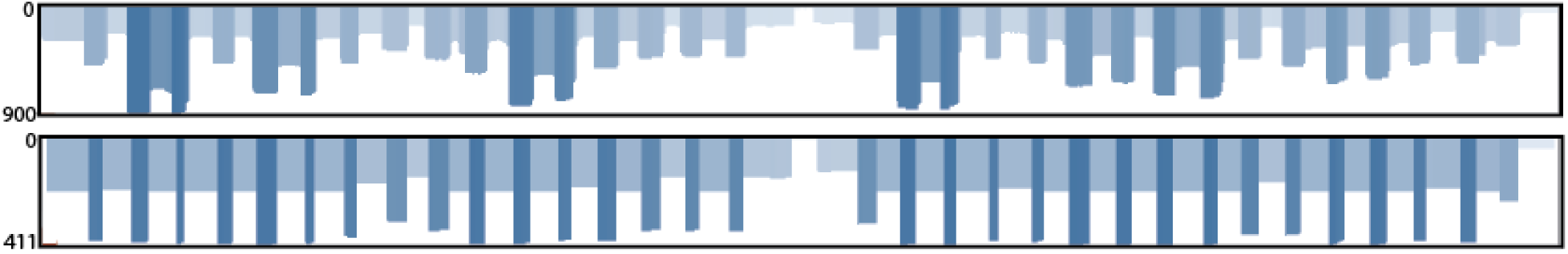
Coverage plots for ZikaAsian scheme sequenced on MinION before (top panel) and after primer trimming and coverage normalisation (bottom panel). During the preprocessing step reads are trimmed using a BED file containing primer positions and read coverage is normalised. The coverage plot was produced using the Tablet genome viewer ^30^.

#### Illumina Sequencing

We compared metagenomic sequencing to the ZikaAsian scheme with the Illumina MiSeq protocol using five clinical samples of Zika from Columbia. Using a previously described method for metagenomics sequencing^2,16^, only a small percentage (< 0.01%) of our reads aligned to Zika virus and they covered only a fraction of the genome (**Table 1** ). Using the ZikaAsian scheme, we were able to generate high coverage of all the genomes (**Table 4**). Illumina sequencing reads are available from BioProject PRJNA358078 (https://www.ncbi.nlm.nih.gov/bioproiect/?term=PRJNA358078).

**Table 4.** Results of amplicon scheme sequencing on five Zika-positive clinical samples from Columbia using the ZikaAsian scheme on the Illumina MiSeq.

| Sample | Ct <sup>1</sup> | GE/ul RNA <sup>2</sup> | Reads | ZIKV reads <sup>3</sup> | % ZIKV | % Coverage | Depth | Genbank ID |
| --- | --- | --- | --- | --- | --- | --- | --- | --- |
| ZC188 | 34 | 36 | 1,114,568 | 760,976 | 68.3 | 98.3 | 29,395 | KY317936 |
| ZC192 | 33.9 | 39 | 1,246,644 | 795,474 | 63.8 | 98.4 | 30,396 | KY317937 |
| ZC199 | 35.9 | 10 | 2,772,457 | 379,064 | 13.7 | 93.4 | 14,848 | KY317938 |
| ZC204 | 33.6 | 48 | 1,065,517 | 751,872 | 70.6 | 99.7 | 29,003 | KY317939 |
| ZC207 | 35.2 | 16 | 939,820 | 506,821 | 53.9 | 96.7 | 19,478 | KY317940 |

(1) qRT-PCR cycle threshold (Ct) value
(2) Genome copy equivalents (GE) per microlitre calculated from Ct value and standard dilutions.
(3) Number of reads aligning to a Zika virus reference genome

#### A Note on Contamination

Cross-contamination is a serious potential problem when working with amplicon sequencing libraries. Contamination risk is minimised by maintaining physical separation between pre- and post-PCR areas, and performing regular decontamination of work surfaces and equipment, e.g. by UV exposure or with 1% sodium hypochlorite solution. Contamination becomes harder to mitigate against as the viral load in a sample reduces. Processing high viral count samples (e.g. post-isolation) can increase the risk of amplicon ‘bleed-over’ into subsequent low viral count samples. When determining how many PCR cycles to use, begin with a lower number and increase gradually to avoid generating excessive quantities of amplicons which makes physical separation harder.

The best safeguard to help detect contamination is the use of negative controls. These controls should be sequenced even if no DNA is detected by quantification or no visible band is present on a gel. Negative control samples should be analysed through the same software pipeline as the other samples. The relative number of reads compared with positive samples give a simple guide to the extent of contamination, and inspection of coverage plots demonstrates can help determine whether a specific region is involved, and potentially its origin by reference to the sequence.

#### Comparison with other approaches

The main benefit to the PCR based approach described here is cost and sensitivity. In theory both PCR and cell culture require only one starting molecule/virion, making them both exquisitely sensitive. In practice, however, the reaction conditions do not allow single-genome amplification and typically a number of starting molecules are required. PCR also has limited sensitivity in cases where the template sequence is divergent from the expected due to primer binding kinetics. This makes it more suitable for use in an outbreak situation where isolates are highly related and low cost per sample and rapid turnaround time are required.

The most similar alternative approach to the one described here is AmpliSeq (Life Technologies). However, this protocol is specific to the Ion Torrent platform and primer schemes must be ordered directly from the manufacturer, and is consequently more expensive per sample. Alternative software to design primer schemes is available, some of which cater specifically for multiplex or tiling amplicon schemes ^18,19,32,33^, and these may perform better when dealing with divergent genomes due to an emphasis on oligonucleotide degeneracy. Primers generated with such software may also be compatible with this protocol, although PCR conditions may require optimisation. In contrast, Primal Scheme is designed with an emphasis on monitoring short-term evolution of known lineages and primer conditions have been optimised for multiplex PCR amplification efficiency.

Propagation with cell culture has been widely used for virus enrichment ^34–36^. This process is time-consuming, requires specialist expertise, and requires high containment laboratories for especially dangerous pathogens. There is also concern that viral passage can introduce mutations not present in the original clinical sample, potentially confounding analysis ^37,38^. In addition culture requires special facilities and skilled personnel and is often unsuccessful.

Oligonucleotide bait probes have also shown promise as an alternative to metagenomics and amplicon sequencing ^39–42^. The method fish for viral nucleic acid sequences by hybridising target-specific biotinylated probes to the DNA sample then pull them down using magnetic streptavidin beads. These methods however are limited by the efficiency of the capture step due to the kinetics of DNA:DNA hybridisations in complex samples, such as the human genome. The complete hybridisation of all probes to targets can take hours (typical protocols suggest a 24 h incubation, although shorter times may be possible) and may never be achieved due to competitive binding by the host DNA. These methods suffer from a coverage bias which worsens at lower viral abundances resulting in increasingly incomplete genomes, as demonstrated by recent work on Zika ^43^. These methods work best on higher abundance samples, and may not have the sensitivity to generate near-complete genomes for the majority of isolates in an outbreak. Probes for hybridisation capture are more expensive than PCR primers because they are usually designed in a fully-overlapping 75 nt scheme which can run to hundreds of probes per virus and thousands for panels of viruses.

## ETHICS STATEMENT

This study has been evaluated and approved by Institutional Review Boards at The Scripps Research Institute and relevant local IRBs in Colombia and Brazil for Zika and chikungunya sample collection and sequencing.

## ACKNOWLEDGEMENTS

The authors would like to thank the Brazilian Ministry of Health and the LACEN’s of Natal, João Pessoa, Recife, Maceió and Salvador for their support. We would like to thank Terry Fredeking from Antibody Systems for providing the Zika virus samples from Colombia. We thank Kirstyn Brunker for testing the Primal Scheme software. The Zika in Brazil Real-time Analysis (ZiBRA) project (zibraproject.org) is supported by the Medical Research Council/Wellcome Trust/Newton Fund Zika Rapid Response Initiative (grant number ZK/16-078) which also supports J.Q.’s salary. N.J.L. is supported by a Medical Research Council Bioinformatics Fellowship as part of the Cloud Infrastructure for Microbial Bioinformatics (CLIMB) project. Primal Scheme is hosted on the CLIMB platform, where pipeline devleopment and MinION data analysis was performed ^44^. N.D.G. is supported by a National Institutes of Health (NIH) training grant 5T32AI007244-33. K.G.A. is a PEW Biomedical Scholar, and his work is supported by an NIH National Center for Advancing Translational Studies Clinical and Translational Science Award UL1TR001114, and National Institute of Allergy and Infectious Diseases (NIAID) contract HHSN272201400048C. A.B. and T.B. were supported by NIH awards R35 GM119774 and U54 GM111274. T.B. is a Pew Biomedical Scholar. A.B. is supported by the National Science Foundation Graduate Research Fellowship Program under Grant No. DGE-1256082. Work at the Paul-Ehrlich-Institut was supported by a grant “Sicherheit von Blut(produkten) und Geweben hinsichtlich der Abwesenheit von Zikaviren” from the German Ministry of Health. This study was supported by USAID Emerging Pandemic Threats Program-2 PREDICT-2 (cooperative agreement AID-OAA-A-14-00102). The contents are the responsibility of the authors and do not necessarily reflect the views of USAID or the US government.

## AUTHOR CONTRIBUTIONS

J.Q. and N.J.L conceived the project. J.Q., N.D.G., K.G.A and N.J.L. designed the experiments and wrote the manuscript. J.Q., A.S. and O.G.P. designed the online primer design tool. J.T.S. modified nanopolish to support R9/R9.4 data and indels. M.L. wrote the demultiplexing software. N.J.L. designed and implemented the MinION bioinformatics pipeline. N.D.G., K.G., G.O., R.R-S. and K.G.A. developed the Illumina sequencing protocol and bioinformatics pipeline. L.L.L-X collected the chikungunya sample, did clinical diagnosis and received local approvals. S.B. and J.B. performed molecular diagnostics and curated Zika and Chikungunya control material. N.D.G, S.T.P., I.M.C., K.G., G.O., R.R-S., T.F.R., N.A.B., J.J.G, M.G., S.H. and A.B. performed the experiments. All other authors tested the protocol and provided feedback. All authors have read and approved the contents of the manuscript.

## SUPPLEMENTARY INFORMATION

**Table 1.**
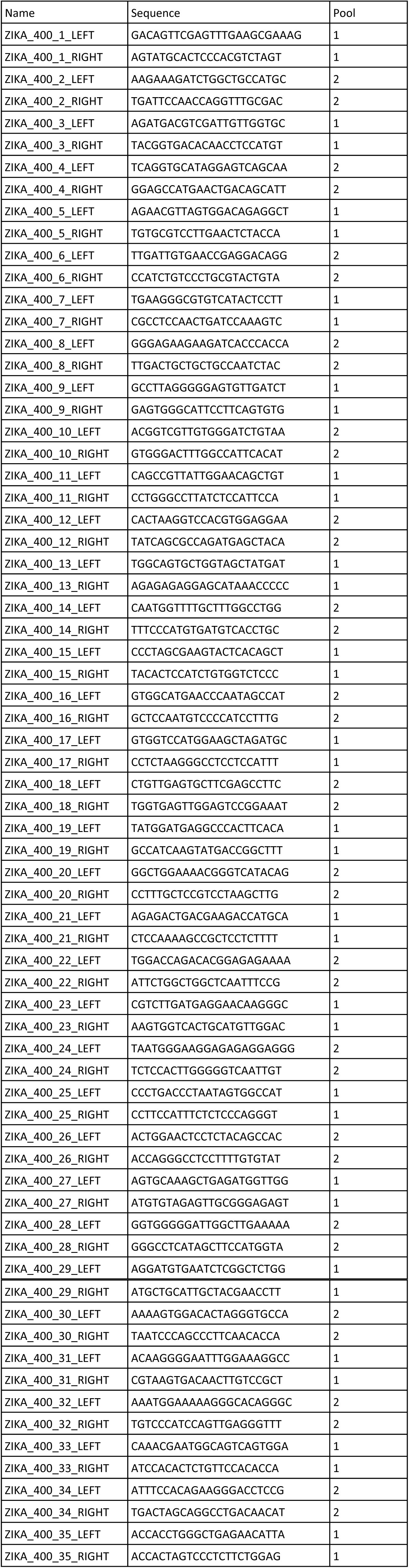
Zika virus “ZikaAsian” scheme used by ZiBRA project generated by the Primal Scheme software ^15^.

**Table 2.**
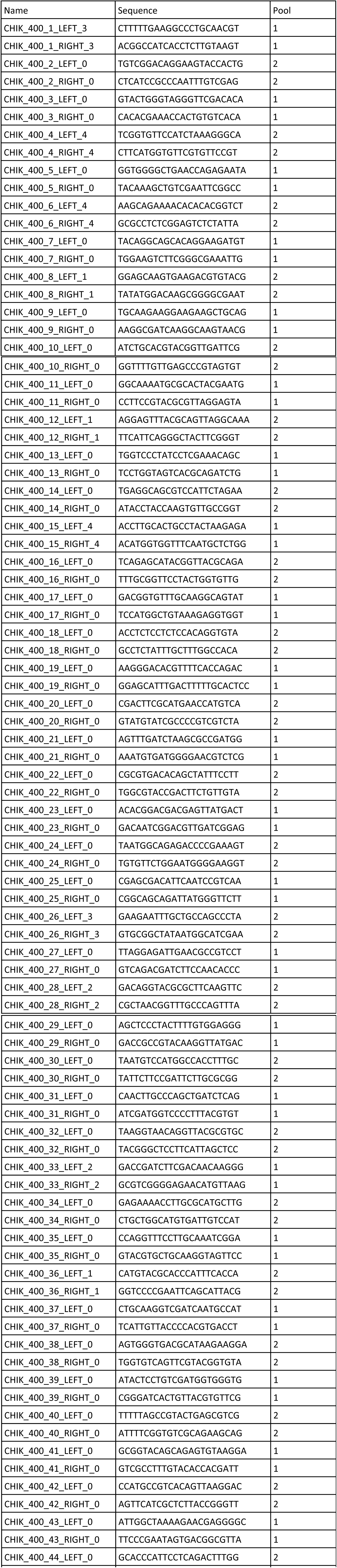
Chikungunya virus “ChikAsiaECSA” 400 nt scheme

